# Cell-Type-Selective Cortical Pathology and Functional Deficits in Synucleinopathy

**DOI:** 10.64898/2026.06.08.730454

**Authors:** Xiaofeng Yang, Changyi Ji, Soomin Song, Nina Harano, Yan Lin, Einar M. Sigurdsson

**Author notes:** Corresponding author: Einar M. Sigurdsson, Department of Neuroscience, Department of Psychiatry, Institute for Translational Neuroscience, New York University Grossman School of Medicine, Science Building, 11th floor, 435 East 30th Street, New York, NY 10016.

## Abstract

Aggregates of α-synuclein (α-syn), a hallmark of synucleinopathies, accumulate in the cerebral cortex accompanied by the emergence of motor symptoms, which are associated with altered cortical neuronal activity. However, the mechanism by which α-syn pathology drives cortical network dysfunction, and how these alterations contribute to impaired motor execution and learning, remain unknown. Here, we adopted a multi-disciplinary approach to elucidate the pathophysiological characteristics in transgenic mice that express mutant human α-syn, with minimal nigrostriatal degeneration. In vivo two-photon imaging revealed distinct alteration patterns in excitatory and parvalbumin (PV)-expressing inhibitory cortical neurons accompanying fine motor deficits during learning. Cell type specific ex vivo whole-cell recording further revealed selectively altered intrinsic properties in excitatory but not PV neurons, consistent with the preferential accumulation of α-syn inclusions in excitatory rather than PV neurons within the same cortical region. These results indicate cell-type selective vulnerability in motor cortex of early stage synucleinopathy, leading to disrupted excitatory/inhibitory balance and dysregulated cortical plasticity, driving early-stage motor symptoms. This study provides evidence for selective vulnerability of excitatory neurons in cortical synucleinopathy.

## Introduction

Misfolding and aggregation of α-synuclein (α-syn) is a defining pathological feature of neurodegenerative disorders collectively known as synucleinopathies. These include Parkinson’s disease (PD), Dementia with Lewy Bodies (DLB) and multiple system atrophy (MSA) [1–3]. In addition, α-syn pathology is also detected in ∼60% of both sporadic and familial cases of Alzheimer’s disease (AD) [4]. These disorders are characterized by progressive and region-specific neuronal loss, accompanied by the accumulation of α-syn aggregates, and are manifested by progressive motor and cognitive deficits [5, 6].

Rather than affecting the nervous system uniformly, α-syn pathology preferentially targets specific neuronal populations, giving rise to distinct clinical phenotypes and progression features. Understanding the basis, pathophysiological mechanisms, and functional consequences of this selective vulnerability remain a central challenge in the field [7]. Many studies have characterized the preferential degeneration of dopaminergic neurons in the substantia nigra pars compacta (SNc) and progressive dopamine depletion in the striatum in synucleinopathy, which underlies the cardinal motor symptoms of Parkinson’s disease. Notably, neighboring dopaminergic neurons in the ventral tegmental area (VTA) are relatively spared, highlighting that even closely related neuronal populations can exhibit markedly different vulnerability profiles, likely due to differences in Ca^2+^ buffering capacity, gene expression level, and connectivity [8–11].

Beyond subcortical dopaminergic systems, increasing evidence points to significant cortical involvement in synucleinopathies [12, 13], particularly in motor cortex, the key region involved in motor execution and learning that is composed of heterogeneous neuronal populations, including excitatory and inhibitory subtypes. Given the precisely balanced cortical network operating near a critical state for efficient information processing and plasticity [14], disruption of this balance may impair normal motor function and potentially accelerate disease progression. Key contributors to maintaining cortical excitatory/inhibitory (E/I) balance include excitatory pyramidal neurons and parvalbumin (PV) interneurons, the latter comprising ∼50% of cortical inhibitory interneurons and playing essential roles in movement initiation [15] and learning [16]. However, whether α-syn pathology selectively impacts cortical subpopulations remains largely unknown.

Recent studies have revealed altered cortical response in somatosensory cortex [12, 17] and visual cortex [18] during sensory perceptions in α-syn mouse models, although conflicting results were reported regarding neuronal activity ranging from hypoactivity to hyperexcitability. Such discrepancies may be attributable to differences in mouse models and region-specific pathophysiological features. In the motor cortex, functional alterations in different types of cortical neurons were also observed in vitro [19] or in vivo [20] under resting conditions, further suggesting that selectively vulnerability of cortical neurons may contribute to motor deficits.

Nevertheless, it remains unclear whether the real-time cortical activity that encodes motor outputs are disrupted in synucleinopathy, how neuronal subtypes are differentially affected, and how altered cortical networks adapt throughout the learning process. Addressing these questions requires direct assessment of cortical population combined with sensitive analyses of the associated fine motor deficits in behaving mice.

In the present study we adopted a multi-disciplinary approach to elucidate the pathophysiological characteristics in transgenic mice (M83) that overexpress the familial A53T mutant human α-syn, with minimal nigrostriatal degeneration [21]. Cell-type specific two-photon imaging was applied targeting layer 2/3 of primary motor cortex (M1) in mice undergoing motor training on a custom-developed forelimb running platform, which integrated precise 3D tracking of movements.

We observed motor deficits and impaired learning in M83 mice, accompanied by distinct alterations in excitatory and PV inhibitory cortical neurons and a lack of learning-induced functional reorganization. Electrophysiological analyses further revealed selectively altered intrinsic firing properties, indicating that disruption of cortical E/I balance, arising from selective neuronal vulnerability to α-syn pathology, drives early-stage motor symptoms.

## Results

### Fine motor deficits and impaired learning in the early stage of synucleinopathy

To assess motor control and learning ability, homozygous M83 mice in the early stages of motor impairment (8–9 months of age) and their age-matched non-transgenic (nTg) controls of the same strain background were trained on a custom-developed motor training platform (**Fig. 1A**). All subject mice underwent 30-minute motor training in 6 consecutive sections, each comprising 5 forced forelimb running trials (**Fig. 1A**). For gait analysis, forelimb movements of head-fixed mice were captured using infrared camera and analyzed with a deep learning-based analyzing tool (DeepLabCut) [22] to extract 3D forepaw positions during running (**Fig. 1B-C**).

**Figure 1.**
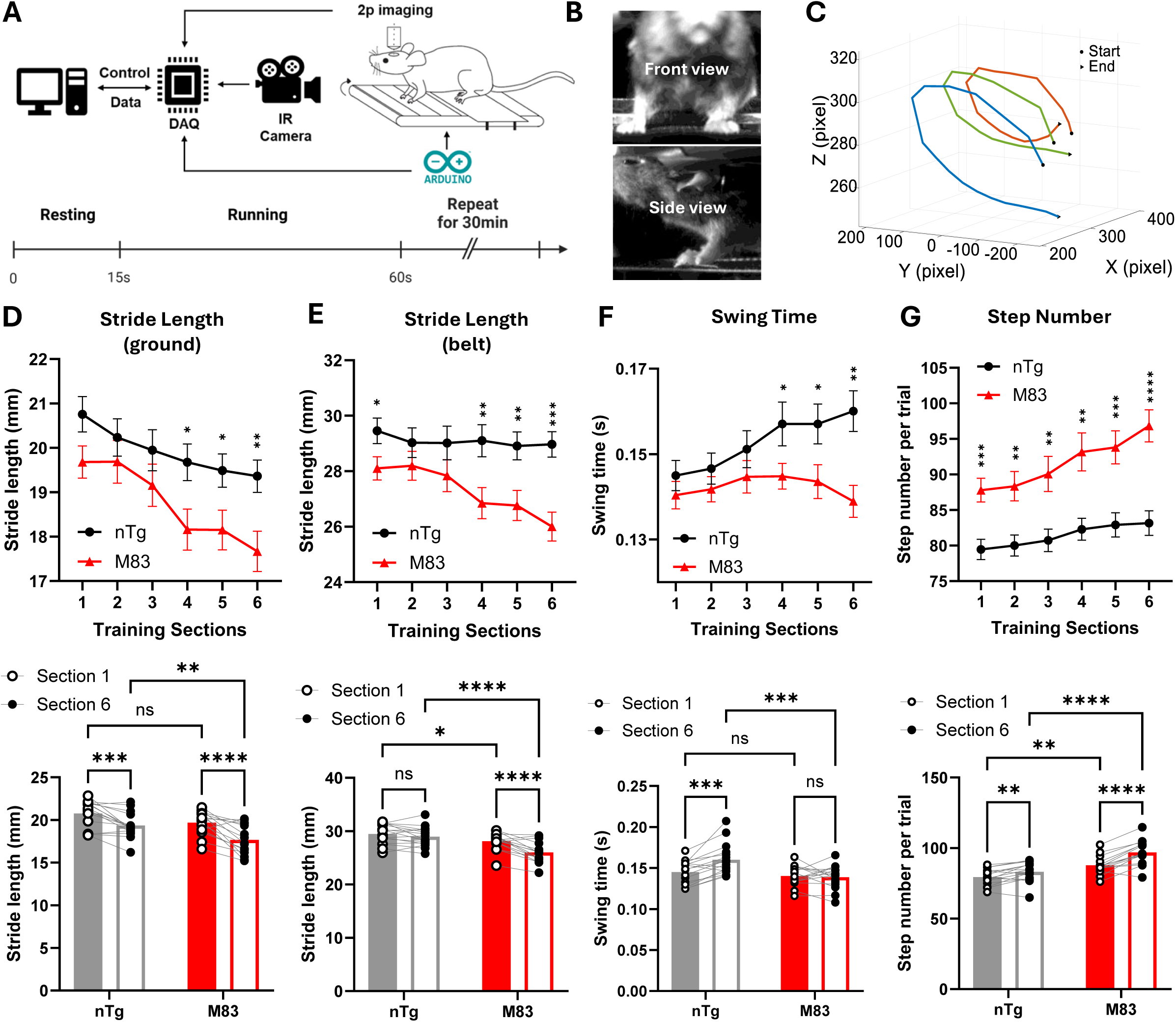
Motor deficits of M83 mice revealed in treadmill running task. **A.** Schematic of treadmill training platform under two-photon microscope. The treadmill task was controlled by a programmable microcontroller and integrated with infrared behavioral camera recordings. Timestamps from the behavioral camera, treadmill controller, and two-photon imaging system were synchronized through a data acquisition (DAQ) card. Each running trial consisted of 13-15 s of resting followed by 45 s of running. **B.** Front and side view of a head-restrained mouse captured during treadmill running. **C.** Movement trajectories of the right forepaw in 3 example steps in the 3D space identified using DeepLabCut. **D.** Stride length relative to the ground shortened in both nTg and M83 mice after training, with M83 mice showing a significantly greater reduction. This parameter calculates the distance between two consecutive landings of the same paw over the stationary surface. Upper graph: mean stride length relative to the ground across all sections of training. Lower graph: statistical comparison between section 1 and section 6 only. Similar layout applies to Figures E–G. **E.** Stride length relative to the running belt shortened only in M83 mice over training, whereas nTg mice maintained a stable stride length. This parameter calculates the distance between two consecutive landings of the same paw in relation to the moving treadmill belt. **F.** Swing time of nTg mice increased significantly after training while this change was absent in M83 mice. This parameter calculates the time spent in the air between two consecutive landings of the same paw. **G.** Step number of M83 mice within the 45s running window was significantly higher than that of nTg mice, and motor training further enhanced this difference. (D-G) nTg n=16 mice, M83 n=15 mice. Two-way ANOVA with Tukey’s multiple comparisons test (all training sections). Two-way ANOVA with Fisher’s LSD multiple comparison test (section 1 vs section 6) *, **, ***, ****: P < 0.05, 0.01, 0.001, 0.0001.

We first quantified the mean stride length of a single paw relative to both the ground and the moving belt across all steps. Stride length relative to the ground decreased in both groups over the 30 min running session, suggesting a shift toward a more upright posture, with M83 mice exhibiting a more pronounced reduction than nTg mice (**Fig. 1D**). When measured relative to the moving belt, nTg mice maintained consistent stride length throughout training, while M83 mice failed to do so, exhibiting a shortening of stride length over repeated running trials (**Fig. 1E**). As stride length is largely determined by the swing time of each step cycle, we next calculated the swing time and found that swing time was comparable between nTg and M83 mice during the initial training sections. With repeated training, nTg mice gradually increased the swing time, which compensated for the reduced stride length relative to the ground and helped maintain stride length relative to the moving belt. However, this progressive adaptation to treadmill running was not observed in M83 mice (**Fig. 1F**), suggests impaired motor learning. As a result of reduced stride length, M83 mice increased the number of steps to cover the same running distance in each trial, whereas nTg mice maintained a relatively stable step number (**Fig. 1G**). The drastically increased step number in M83 mice suggests an inherent consequence of altered gait patterns and could serve as a strategy that M83 mice developed to compensate for their motor deficits.

During the acquisition of novel motor skills, movement variability across repeated trials is a key feature, typically observed in the early stage of learning, which is believed to facilitate motor exploration and is subsequently modulated by outcomes into gradually refined movement patterns [23]. To assess this kinematic feature of learning, we quantified the pairwise correlation of 3D forelimb trajectories across steps in each training section. We observed a significant increase in step-to-step correlation of movement trajectories in nTg mice, indicating a gradual reduction in motor variability over learning (**Fig. 2A, C, D**). Interestingly, M83 mice exhibited significantly higher correlations than nTg mice even at initial trials, and this high-level correlation persisted throughout training (**Fig. 1B, C, D**). Given that the task was novel for both genotypes, these findings suggest abnormally decreased variability of forelimb movements in M83 mice, indicating reduced motor flexibility and an early sign of rigidity or stiffness, one of the cardinal motor symptoms that can already be observed during the early stages of PD [24].

**Figure 2.**
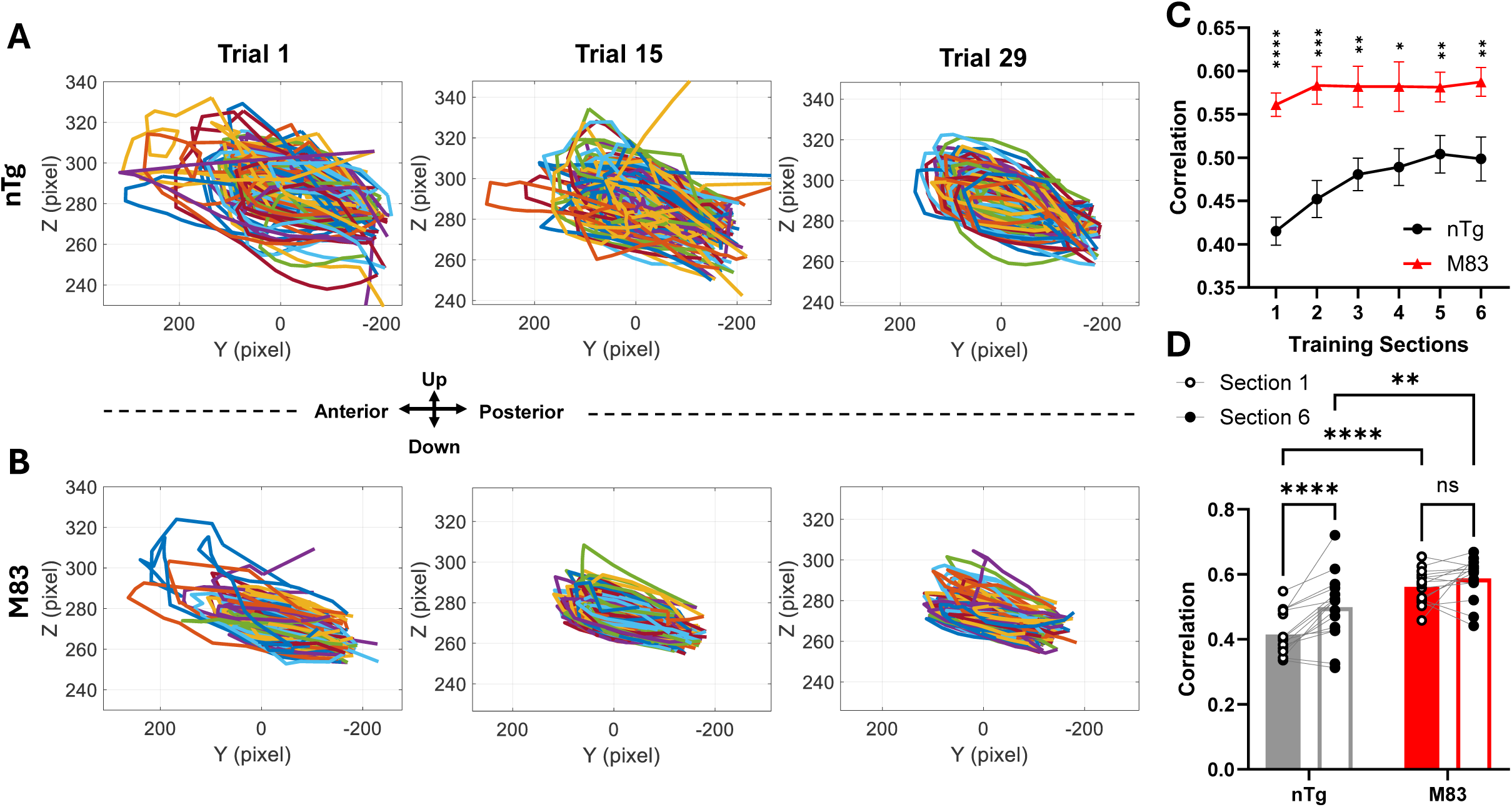
Cross-step correlation analysis revealed altered movement kinematics in M83 mice. **A.** Single forepaw trajectories of a representative nTg mouse from the side view of all the steps during early, middle, and late training trials. Different colors represent individual steps. nTg mice exhibited more variable trajectories during early trial, reflecting greater exploratory behavior in the novel motor task, which gradually became more refined and converged into more stereotyped, repeatable patterns with training. **B.** Single forepaw trajectories of a representative M83 mouse from the side view of all the steps during early, middle, and late training trials. Compared to nTg mice, M83 mice already exhibited less variable step trajectories in trial 1, which remain relatively constrained across training. **C-D**. Trial-averaged pairwise correlation of 3D trajectories across all steps in different training sections. M83 mice displayed significantly higher and stable correlation compared to nTg mice, while nTg mice showed a learning-induced gradual increase in correlation over training. (C-D) nTg n=16 mice, M83 n=15 mice. Two-way ANOVA with Tukey’s multiple comparisons test (all training sections). Two-way ANOVA with Fisher’s LSD multiple comparison test (section 1 vs section 6) *, **, ***, ****: P < 0.05, 0.01, 0.001, 0.0001.

In summary, through fine motor assessment, this study provides experimental evidence of subtle motor deficits at the early stage of synucleinopathy and reveals impaired motor learning accompanied by rigidity-like features. These behavioral findings facilitate further investigations of the underlying neuronal pathophysiology in this mouse model.

### Selectively altered Ca^2+^ activity in the motor cortex accompanied by early-stage motor deficits in M83 mice

To characterize the functional properties of excitatory and inhibitory PV neurons in the motor cortex during the forelimb running, we injected AAVs driven by CaMKII or S5E2 promoters [25] to selectively express the Ca²⁺ indicator GCaMP in excitatory neurons or PV interneurons in layer 2/3 of M1, a key region in motor learning [26, 27] (**Fig. 3A**). Populational cortical activities were recorded during the whole training process by two-photon microscope for their responsive features and evolving patterns (**Fig. 3B**).

**Figure 3.**
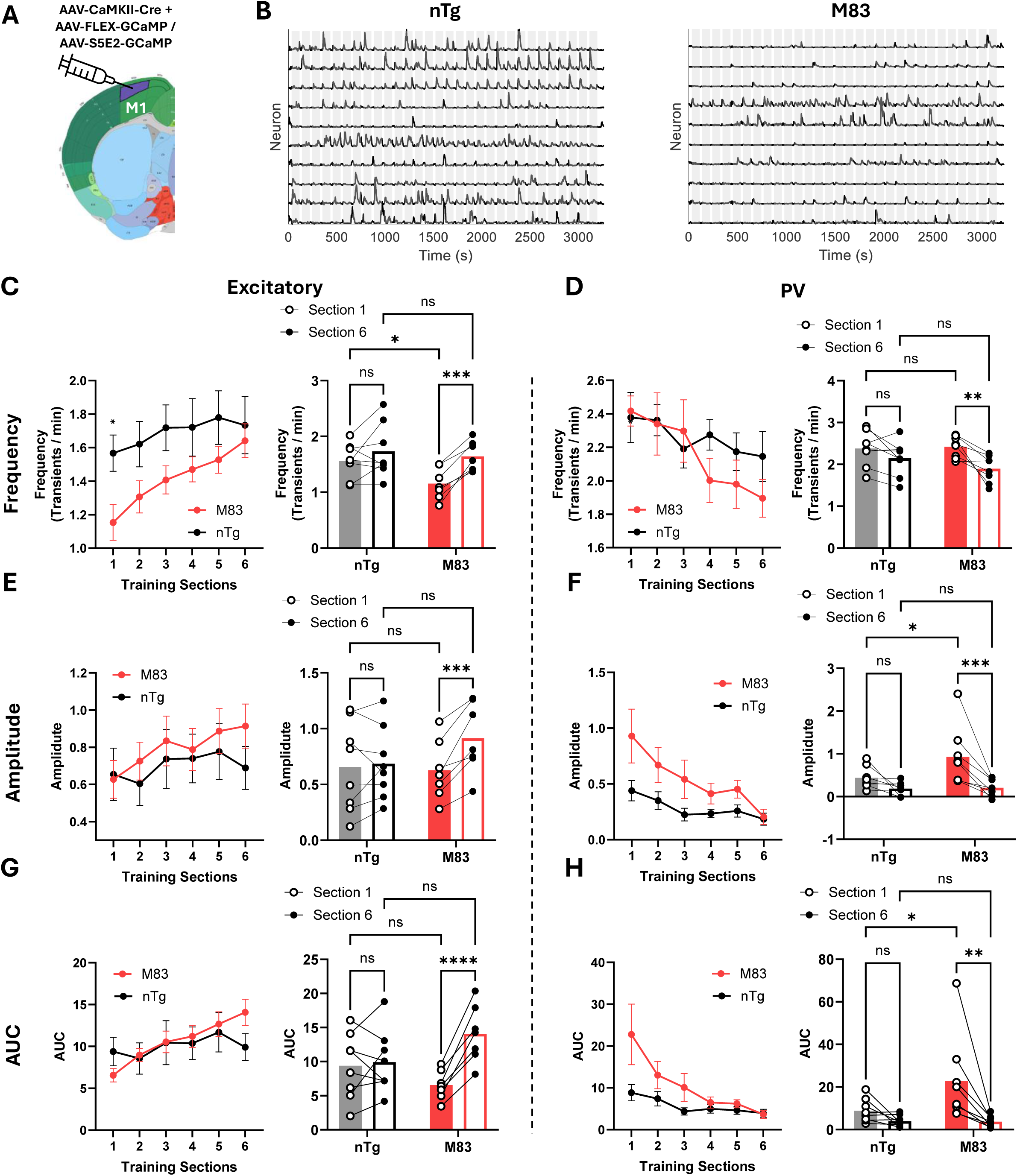
Altered cortical activity in the M1 of M83 mice during running. **A.** AAV viral vectors were injected into layer 2/3 of the primary motor cortex (M1) for Ca^2+^ imaging of excitatory neurons or PV interneurons. Cell-type–specific expression of GCaMP was achieved using the CaMKII promoter for excitatory neurons and the S5E2 promoter for PV interneurons. **B.** Example Ca^2+^ activity traces from 10 excitatory cortical neurons in representative nTg and M83 mice during motor training. Grey bands indicate running periods. nTg mice exhibited more frequent Ca^2+^ events compared with M83 mice. **C-D**. Mean frequency (transients/min) of Ca^2+^ transients of excitatory and PV neurons during forelimb running in nTg and M83 mice. In M83 mice, significantly reduced frequency was observed in the excitatory neurons but not in PV neurons in early section. Through repeated practice, excitatory and PV neurons in M83 mice exhibited significant, gradual changes in activity frequency in opposite directions, whereas nTg mice showed stable activity frequencies across training sections in both cell types. **E-F**. Mean amplitude of Ca^2+^ transients of excitatory and PV neurons during forelimb running for nTg and M83 mice. In M83 mice, significantly increased amplitude was observed in the PV neurons but not in excitatory neurons in early section. Like the evolving trend observed in frequency, M83 mice exhibited significant and opposing gradual changes in Ca^2+^ transient amplitude between excitatory and PV neurons, whereas nTg mice showed stable amplitude across training sections in both cell types. **G-H**. Integrated area under curve (AUC) of Ca^2+^ activity during the running period of excitatory and PV neurons for nTg and M83 mice. In M83 mice, significantly increased AUC was observed in the PV neurons but not in excitatory neurons in early section. Like the evolving trend observed in frequency and amplitude, M83 mice exhibited significant and opposing gradual changes in AUC between excitatory and PV neurons, whereas in nTg mice this parameter maintained stable across training sections in both cell types. (C-H) nTg excitatory n=8 mice, M83 excitatory n=7 mice, nTg PV n=8 mice, M83 PV n =8 mice. Two-way ANOVA with Tukey’s multiple comparisons test (all training sections). Two-way ANOVA with Fisher’s LSD multiple comparison test (section 1 vs section 6) *, **, ***, ****: P < 0.05, 0.01, 0.001, 0.0001.

At the naïve stage of learning (section 1), excitatory neurons in M83 mice exhibited a significantly reduced frequency of Ca^2+^ transients compared to nTg mice during running (**Fig. 3C**). In contrast, the activity of PV inhibitory neurons was comparable between the two groups, indicating a hypoactive cortical state of M83 mice (**Fig. 3D**). Notably, with continued practice, excitatory neuronal activity in M83 mice gradually increased, accompanied by an opposite decrease in PV neuron activity, whereas the activity levels for both cell types remained stable in nTg controls (**Fig. 3C-D**).

Furthermore, the amplitude of Ca^2+^ transients also exhibited an opposite evolving pattern during the learning process in M83 mice, between excitatory and PV neurons (**Fig. 3E-F**). Notably, the amplitude of Ca^2+^ events in PV neurons was significantly higher in M83 mice at naïve stage of learning (**Fig. 3F**), which suggests enhanced spike frequency in PV neurons, as Ca^2+^ transient amplitude can be approximated by the convolution of the underlying spike trains [28, 29]. A similar phenomenon was observed for the integrated area under the curve (AUC), which reflects the cumulative Ca^2+^ response, incorporating Ca^2+^ influx, intracellular Ca^2+^ release, and signal decay kinetics, rather than peak amplitude alone (**Fig. 3G-H**). These findings indicate cell-type-selective functional alterations and a disrupted cortical E/I balance in M83 mice, characterized by a shift toward network suppression, which is gradually reshaped through motor practice.

As the principal cell type in layer 2/3 of motor cortex, excitatory neurons are responsible for sensory integration and descending motor commands. Hypoactive excitatory neurons could profoundly impair normal motor function. To further examine the distribution of hypoactive neurons within the excitatory population and the training-induced changes, we categorized excitatory neurons into hyperactive, normal, and hypoactive groups in reference to our previous work and related studies [30, 31] (**Fig. 4A-B**). Although, the fraction of hypoactive neurons in M83 mice was significantly higher than in nTg mice at section 1, it gradually decreased to a similar level with nTg mice after training, whereas no change was observed in nTg mice (**Fig. 4C-D**). Neuron ID tracking confirmed that a larger fraction of neurons transitioned from hypoactive to normal in M83 mice than in the opposite direction. In contrast, this exchange was balanced in nTg neurons (**Fig. 4E-F**).

**Figure 4.**
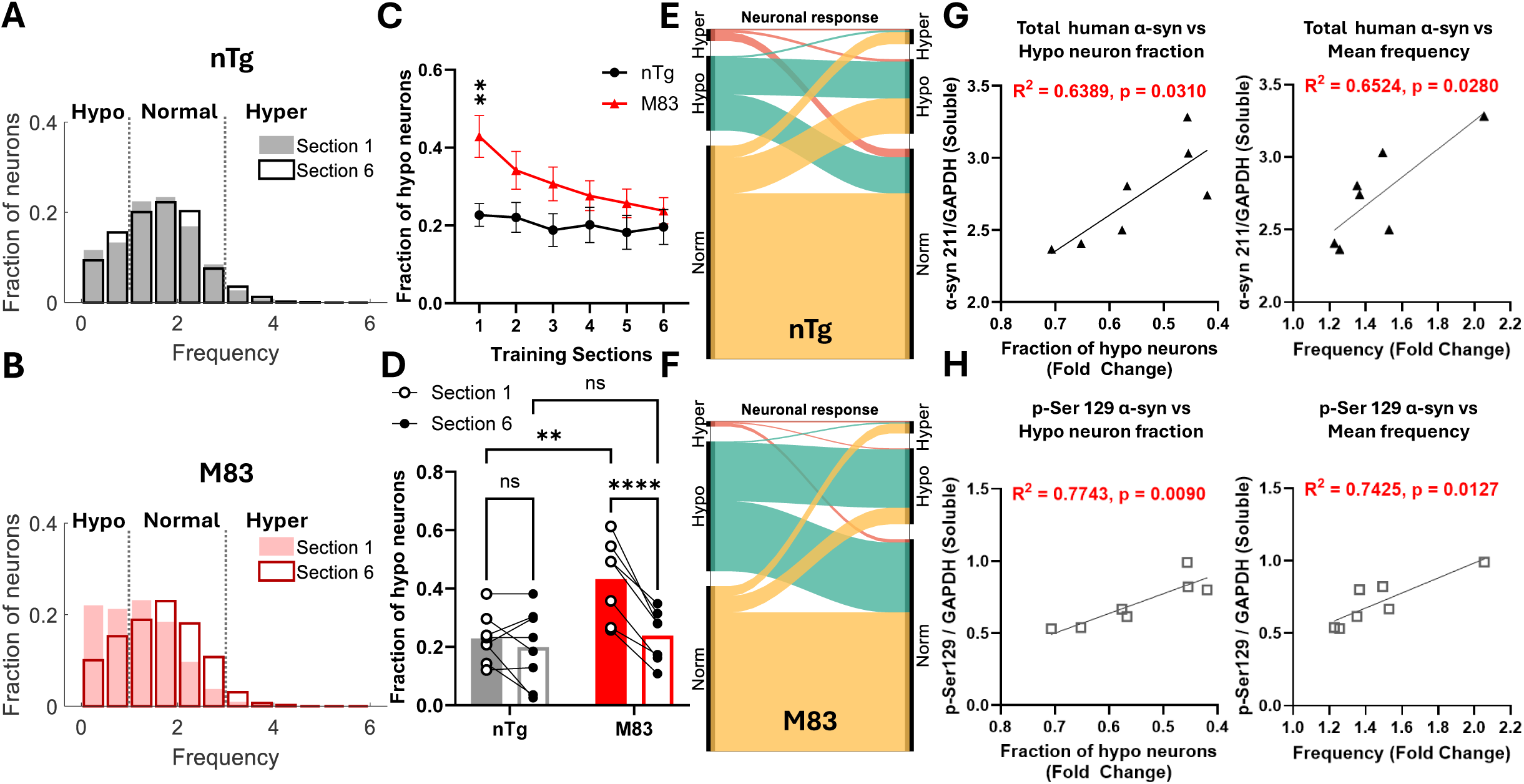
Hypoactivity of excitatory cortical neurons is correlated with pathological α-syn levels in M83 mice. **A-B.** The distribution of all recorded excitatory neurons in nTg (A) and M83 (B) mice in early and late sections of training. Neurons were classified as hypoactive, normal and hyperactive according to their mean frequency during running. Clear change in frequency distribution was demonstrated in M83 mice only. **C-D**. Mean proportion of hypoactive excitatory neurons across different sections of motor training. A larger fraction of hypoactive excitatory neurons was observed in M83 mice during section 1, followed by a gradual return to the levels comparable to nTg mice. **E-F**. The flowchart demonstrates the exchange of neurons across clusters from section 1 (left side) to section 6 (right side). Red: hyperactive neurons in section 1; Green: hypoactive neurons in section 1; Yellow: normally active neurons in section 1. **G-H**. Pathological α-syn levels correlated with changes in Ca^2+^ activity during the training process. The levels of total soluble human α-syn (G) and phosphorylated α-syn (H) were measured by western blot assay identified by α-syn211 and pSer129 antibodies respectively. They were strongly correlated with the fold changes of activity frequency or fraction of hypoactive neurons recorded in vivo during the training process. **(C-D)** nTg excitatory n=8 mice, M83 excitatory n=7 mice. Two-way ANOVA with Tukey’s multiple comparisons test (all training sections). Two-way ANOVA with Fisher’s LSD multiple comparison test (section 1 vs section 6) **, ****: P < 0.01, 0.0001.

To determine whether pathogenic burden of α-syn contributes to the degree of altered cortical activity, we quantified total α-syn levels in mice brain after two-photon imaging through western blotting and assessed their correlation with training-induced activity changes. We found the level of mutant human α-syn or phosphorylated α-syn in soluble fraction strongly correlated with the fold changes of Ca^2+^ transient frequency and the hypoactive fraction of excitatory neurons (**Fig. 4G-H**). These data indicate that pathogenic burden of α-syn proportionally contributes to the extent of hypoactive deviation in cortical neurons.

### Altered temporal dynamics and impaired plasticity in cortical neurons in M83 mice

Although motor practice progressively ameliorated the disrupted E/I balance, motor performance did not improve in M83 mice. We therefore investigated the temporal dynamics of cortical neuronal populations during running. When aligned to running onset, both excitatory and PV neurons in M83 mice exhibited distinct temporal response patterns compared with nTg controls. In nTg mice, excitatory and PV neurons exhibited coherent responses with activity increasing at running onset (**Fig. 5A-B**). Considering the strong recurrent connectivity between excitatory and PV neurons within the cortex [32], as well as recent findings of correlated synaptic strength in reciprocally connected excitatory–PV neuronal pairs [33], this coordinated dynamic is expected. In contrast, M83 mice displayed reduced excitatory activity together with a rapid and pronounced increase in PV inhibitory activity (**Fig. 5C-D**). Notably, this temporal disruption of E/I balance was not rescued by training, potentially contributing to motor deficits and impaired learning.

**Figure 5.**
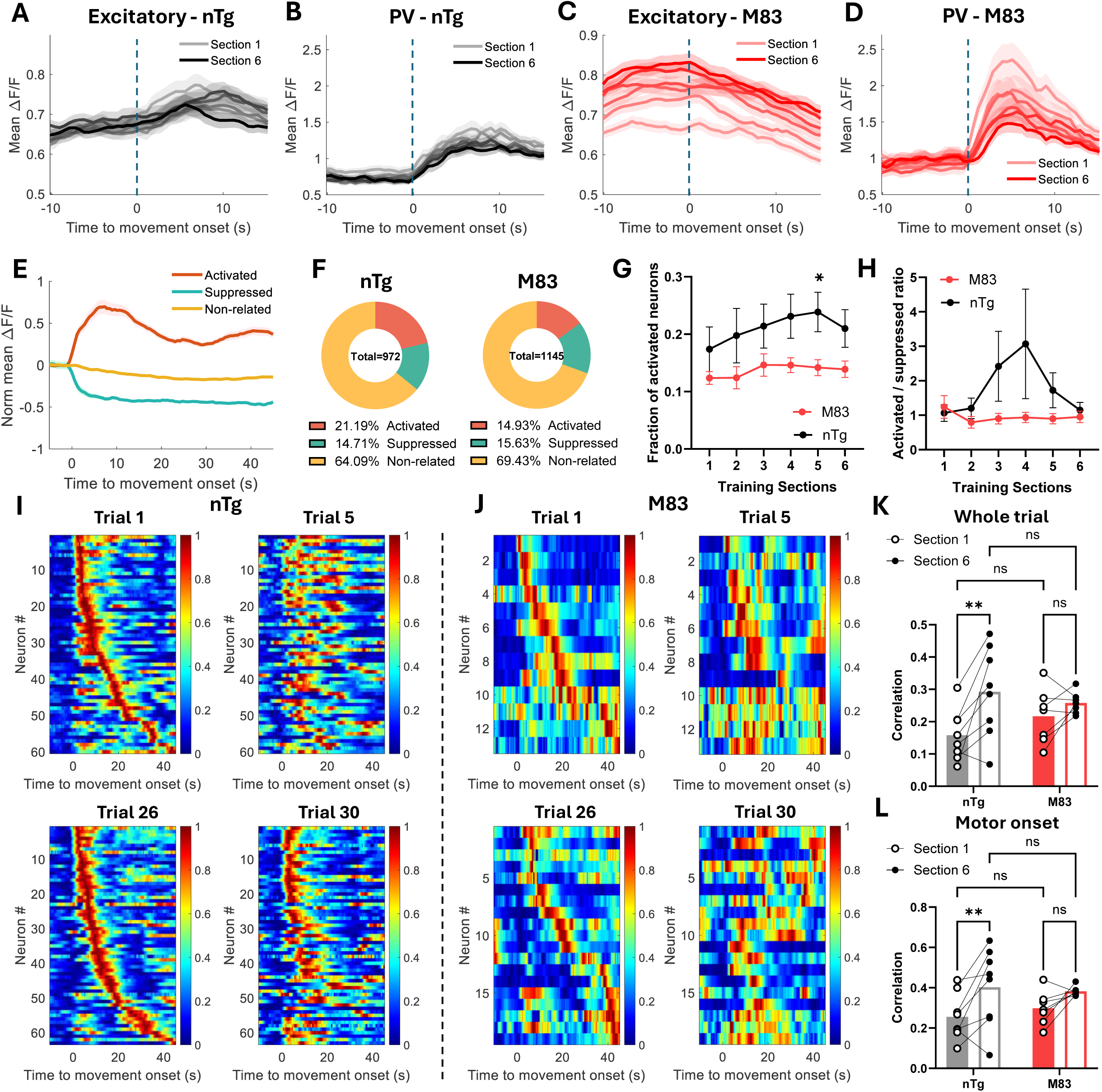
Altered temporal and populational response of cortical excitatory neurons in M83 mice. **A-D.** Trial-averaged temporal Ca^2+^ activity of excitatory and PV neurons in all training sections aligned to movement onset. Color gradients from light to dark marks training section 1 to 6 for nTg (black) and M83 (red) mice. Coherent ramping activity at running onset was observed in the nTg group between excitatory and PV neurons, with stable response levels across training sections. In contrast, the M83 group showed an opposite temporal response between the two cell types and gradual changes in activity levels over training process. Dash line and 0 time point indicate treadmill onset. **E**. Excitatory neurons were classified according to their response preference during running. Ca^2+^ response of subgroups of excitatory neurons in M83 mice during section 1. **F**. Pie chart shows larger fraction of activated neurons in the middle stage of training (section 4) in nTg mice than in M83 mice. **G-H**. Fraction of running activated excitatory neurons, and the ratio between activated and suppressed neurons. The proportion of activated excitatory neurons increased during the middle stage of training, whereas this temporal change was absent in M83 mice. **I-J**. Single-trial population activity of activated excitatory neurons from representative nTg and M83 mice. Firstly, activated neurons were identified according to their response in different training sections: section 1 (trial 1 - 5), section 6 (trial 26 - 30). Then, the same group of identified neurons were sorted according to their peak response time in the first trial of that section (e.g. trial 1 or trial 26). **K-L**. Correlation of response sequence of activated excitatory neurons across all trials within section 1 or section 6. Increased cross-trial correlation in nTg mice indicates plastic changes and the gradual formation of repeatable activity patterns through learning, while this populational adaptation was absent in M83 mice. (G-H, K-L) nTg excitatory n=8 mice, M83 excitatory n=7 mice. Two-way ANOVA with Tukey’s multiple comparisons test (all training sections). Two-way ANOVA with Fisher’s LSD multiple comparison test (section 1 vs section 6) *, **: P < 0.05, 0.01.

To examine plastic changes at the population level in M83 mice, we first classified excitatory neurons based on their response preference to running (**Fig. 5E**). This analysis revealed a reduced fraction of positively responsive neurons in M83 mice compared to nTg controls (**Fig. 5F**). Furthermore, in nTg mice, we observed a temporal increase in the proportion of positively responsive neurons during the middle phase of learning (**Fig. 5G-H**), suggesting a process of exploration and selection in cortical encoding prior to refinement—a phenomenon commonly reported in the motor cortex during skill acquisition [26, 34]. Previous studies showed that early motor learning involves variable population activity that gradually evolves into highly reproducible spatiotemporal sequences, particularly within movement-related neuronal subpopulations [26, 35]. In line with these findings, we detected a progressive stabilization of activity sequences of positively responsive neurons across training in nTg mice (**Fig. 5I**). Specifically, the sequential activation of positively responsive neurons became increasingly repeatable, with reduced trial-to-trial variability after learning compared to the initial stage (**Fig. 5K-L**). In contrast, this learning-induced stabilization was absent in M83 mice (**Fig. 5J**), in which cross-trial correlations of activity sequence remained at intermediate levels throughout learning (**Fig. 5K-L**).

According to previous studies using the same mouse model, pathological α-syn aggregates are absent in immunohistology of dopaminergic neurons of the substantia nigra at this age [21, 36]. Therefore, the observed impaired cortical plasticity is less likely to be driven by dopaminergic degeneration. To further assess potential dopaminergic involvement, we performed stereological quantification of dopaminergic neurons in both the substantia nigra pars compacta (SNc) and ventral tegmental area (VTA), two major midbrain sources of dopamine that are known to modulate plasticity in motor learning [37]. We found comparable numbers of TH-positive neurons in both brain regions of M83 mice compared with age-matched nTg controls (**Fig. 6**).

**Figure 6.**
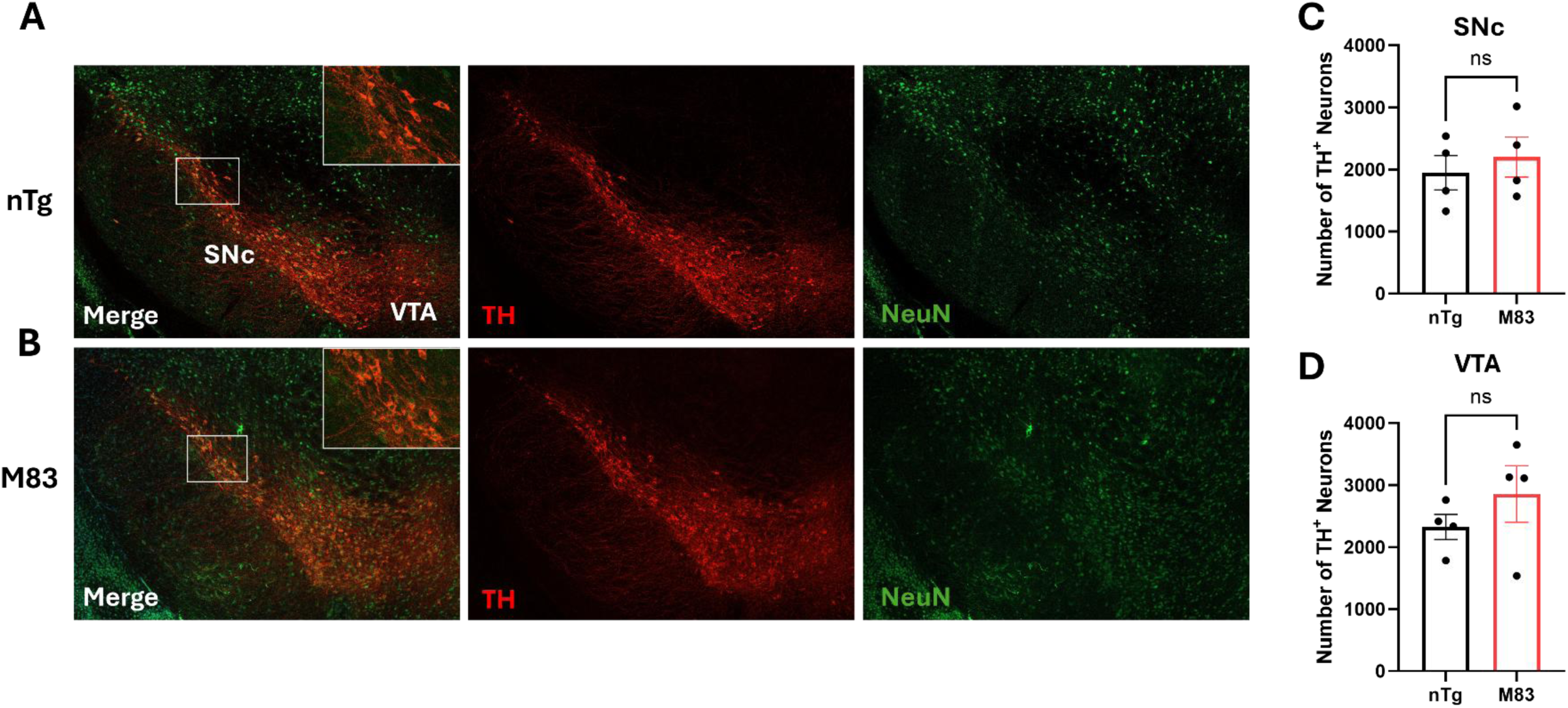
Preserved dopaminergic (DA) neurons in the midbrain of M83 mice. **A**. Representative immunostaining of tyrosine hydroxylase (TH) positive neurons in substantia nigra pars compacta (SNc) and ventral tegmental area (VTA) of 8–9-month female M83 and age matched nTg mice. Left to right: Merged channels, TH staining (Red) and NeuN staining (Green). **B-C**. Stereological count of DA neurons in SNc and VTA from one hemisphere. A series of coronal sections with 210 μm intervals were collected from bregma −2.5 mm to −3.4 mm. No significant difference in TH labelled cells was detected between M83 and nTg control mice, suggesting the absence of nigrostriatal neuron loss in M83 mice at this age. TH positive neurons were counted through ImageJ. nTg n= 4 mice, M83 n= 4 mice. Unpaired Mann-Whitney t-test.

Together, these findings indicate a temporally dysregulated E/I balance in M83 mice throughout learning with reduced plasticity at population level which is necessary for acquiring new motor skills. The PV-neuron-driven suppression may further dampen excitatory recruitment, limiting the engagement of movement-related neuronal populations. As a result, cortical dynamics in M83 mice appear constrained, with reduced flexibility in population responses that may hinder the adaptive reorganization required for efficient motor learning.

### Selectively altered excitability and synaptic transmissions of cortical neurons in M83 mice

As in vivo recordings has revealed distinct alterations in different cell types during running, we next determined whether these dysfunctions arise from intrinsic deficits driven by α-syn pathology. With the guidance of cell-type-specific GCaMP expressions, we performed ex vivo patch-clamp recordings to assess the intrinsic firing properties and synaptic transmissions of layer 2/3 cortical neurons. Neuron excitability was measured by whole-cell current-clamp recording with increasing current injections for their action potential frequency, rheobase current, fast-afterhyperpolarization (fAHP), spike frequency adaptation and dynamic properties [30]. Excitatory neurons exhibited significantly reduced firing frequency in M83 mice compared with nTg mice (**Fig. 7A-B**), accompanied by an increase in rheobase current (**Fig. 7C**), suggesting significantly reduced excitability which may contribute to their hypoactive state observed in vivo, whereas the excitability of PV neurons remained unchanged between the two mouse groups (**Fig. 7F-H**).

**Figure 7.**
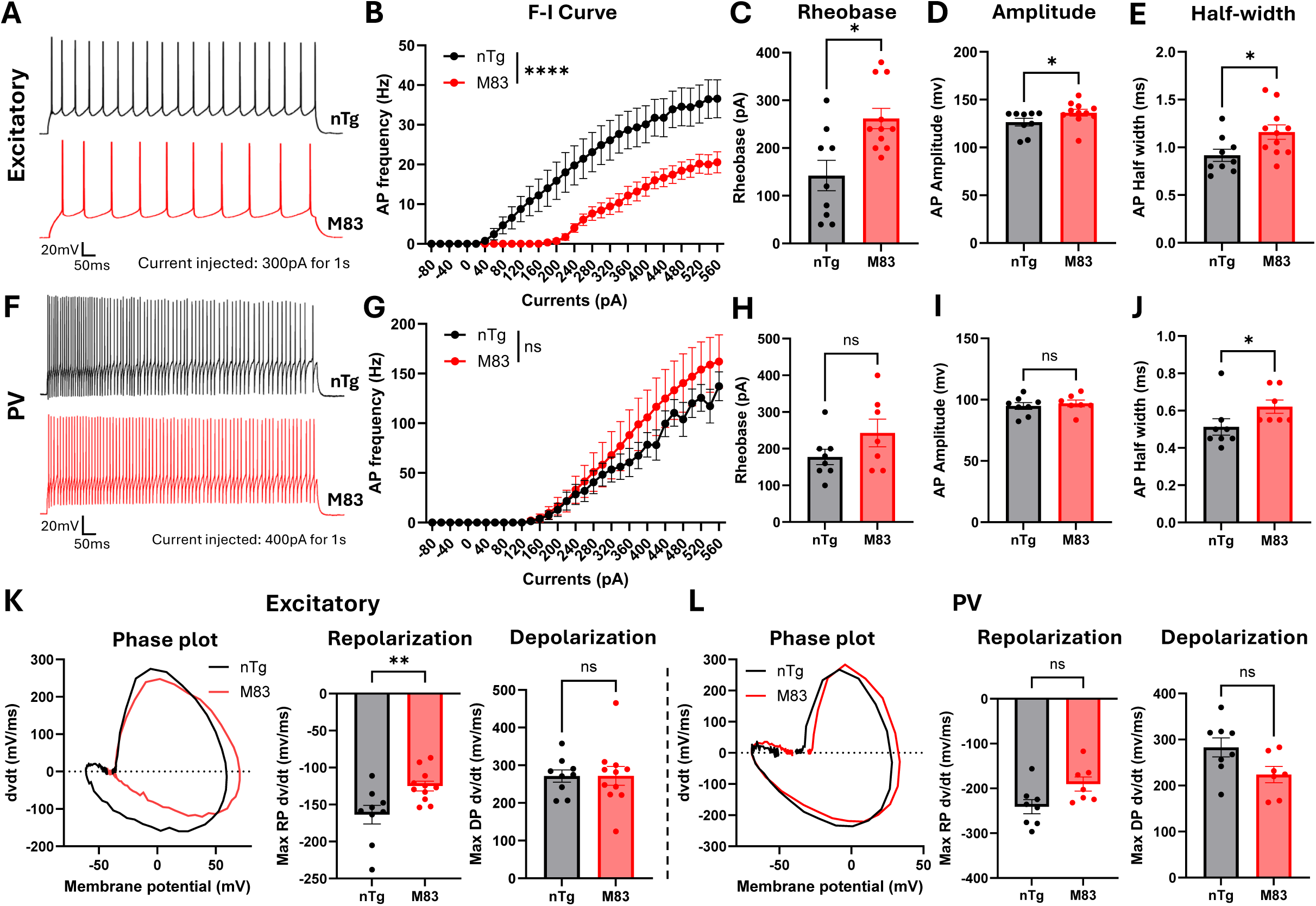
Preferentially reduced excitability of cortical excitatory neurons in M83 mice. **A**. Example traces of current-clamp recording of excitatory neurons in mouse brain slices revealed reduced firing frequency in M83 mice compared to nTg mice. Example traces show neuronal responses to a 300pA depolarizing current injection for 1 s **B**. The F/I curve of excitatory neurons in nTg or M83 mice, recorded during stepwise-increasing current injections. Significantly diminished excitability of cortical excitatory neurons was observed in M83 mice compared with nTg mice. **C-E**. Rheobase current, action potential amplitude and half-width at minimal current injection to trigger an action potential in excitatory neurons were significantly increased in M83 mice compared to nTg mice. **F-J**. The same current-clamp recording analysis in PV neurons. Example traces were recorded with 400pA depolarizing current for 1 s. The excitability of PV inhibitory neurons was not significantly affected. Action potential amplitude showed no difference between M83 and nTg mice, whereas action potential half-width was increased in M83 mice. **K**. Representative phase-plane plots and quantification of maximum depolarization and repolarization rates of action potentials at minimal current injection in excitatory neurons. The maximum repolarization rate significantly decreased in M83 mice compared to nTg mice. **L**. The same kinematic analysis of action potentials at minimal current injection in PV neurons. No significant differences were found between M83 and nTg mice. Excitatory neurons: nTg n=9, M83 n=11; PV neurons: nTg n=8, M83 n=7; Two-way ANOVA with Tukey test; Mann-Whitney test. *, **, ****: p<0.05, 0.01, 0.0001.

Kinematically, both increased action potential amplitude and half-width observed in excitatory neurons of M83 mice suggest slower spike kinetics (**Fig. 7D-E**). But this alteration was less obvious in PV neurons (**Fig. 7I-J**), indicating dysfunction of voltage-gated ion channels. Phase-plane analysis revealed reduced maximum repolarization rate but not maximum depolarization rate in excitatory neurons (**Fig. 7K**), which suggests potentially reduced potassium channel conductance and/or increased sodium channel function. In contrast, the dynamic feature of PV neurons was not significantly altered (**Fig. 7L**). Abnormal properties of fAHP were observed in excitatory neurons of M83 mice (**Fig. 8A-C**), leading to stronger spike-frequency adaptation (**Fig. 8D-E**), while these changes were absent in PV interneurons (**Fig. 8F-J**). These alterations could impair the ability of sustained firing, resulting in a reduction of firing rates.

**Figure 8.**
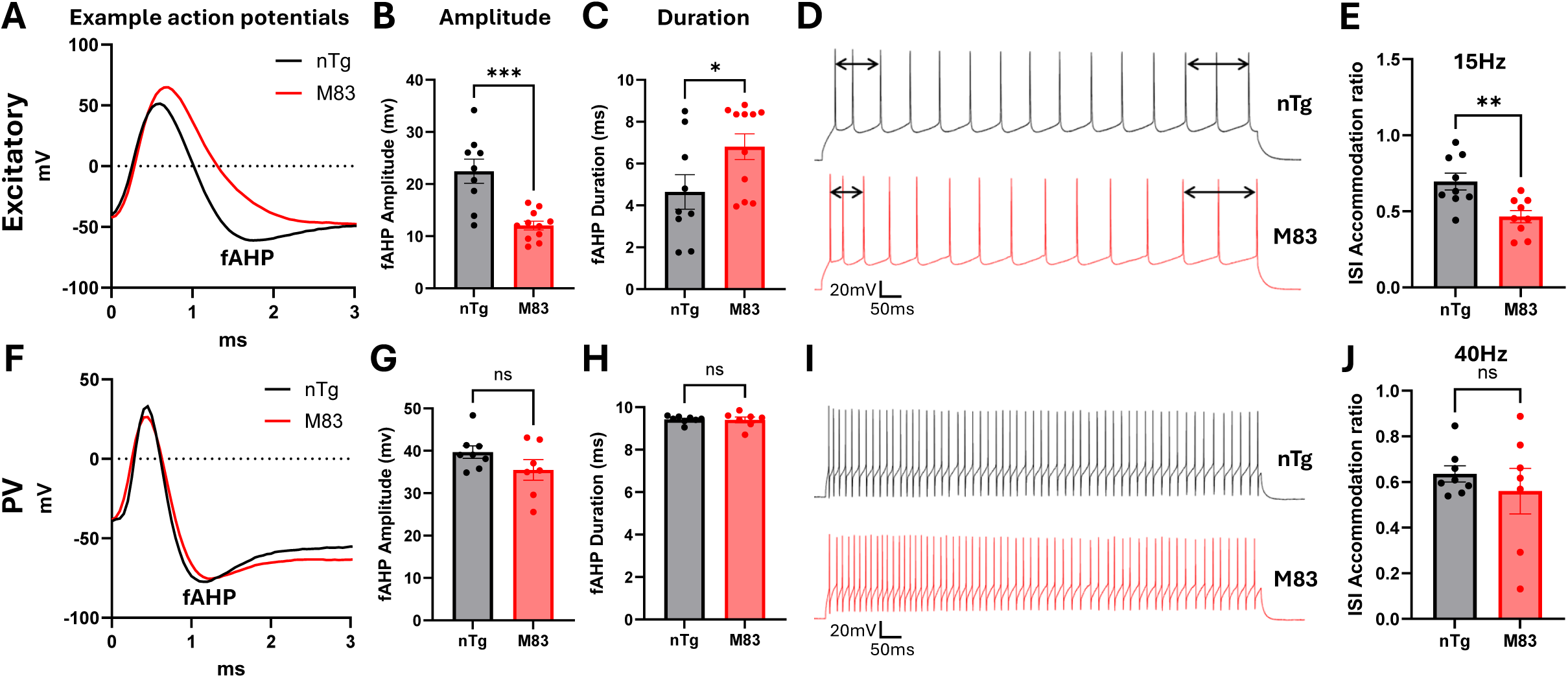
Impaired repetitive firing ability of excitatory cortical neurons in M83 mice. **A**. Representative waveforms of action potentials of excitatory neurons at minimal current injection. Different dynamics were observed between neurons from M83 and nTg mice. **B-C**. Amplitude and duration of the fast afterhyperpolarization (fAHP) were measured in excitatory neurons. A reduced fAHP amplitude and prolonged fAHP duration were observed in M83 mice. **D-E**. Representative recordings of repetitive firing and quantification of firing rate adaptation in excitatory neurons at the minimum current required to evoke firing rates greater than 15 Hz. A smaller inter-spike interval (ISI) accommodation ratio was observed in the M83 group compared to the nTg group, indicating stronger spike probability adaptation (SPA) in M83 mice. **F-J**. The same measurement of repetitive firing ability in PV neurons. The ISI accommodation ratio was quantified at the minimum current required to evoke firing rates greater than 40 Hz. Similar levels of SPA were observed in M83 and nTg groups. Excitatory neurons: nTg n=9, M83 n=11; PV neurons: nTg n=8, M83 n=7; Two-way ANOVA with Tukey test; Mann-Whitney test. *, **, ***: p<0.05, 0.01, 0.001.

Furthermore, we evaluated synaptic transmission in the more vulnerable excitatory neurons through measuring spontaneous excitatory post-synaptic currents (sEPSCs) and spontaneous inhibitory post-synaptic currents (sEPSCs) of the same neuron (**Fig. 9A-B**). Ratio between sEPSCs and sIPSCs can reflect E/I balance at single neuron level [38]. Layer 2/3 excitatory neurons in M83 mice displayed significantly decreased sEPSC and increased sIPSC frequency (**Fig. 9C-D**), whereas the amplitudes for both sEPSC and sIPSC were comparable between M83 and nTg mice (**Fig. 9E-F**). For the same excitatory neuron that was recorded, the E/I ratio of PSCs frequency was significantly decreased in M83 mice compared to nTg mice, but the E/I ratio of PSCs amplitude was preserved across genotypes (**Fig. 9G-H**). These findings further reveal impaired E/I balance at single neurons and synaptic level. Specifically, it suggests an imbalanced synaptic input rather than a change of postsynaptic receptor function at early stage of synucleinopathy.

**Figure 9.**
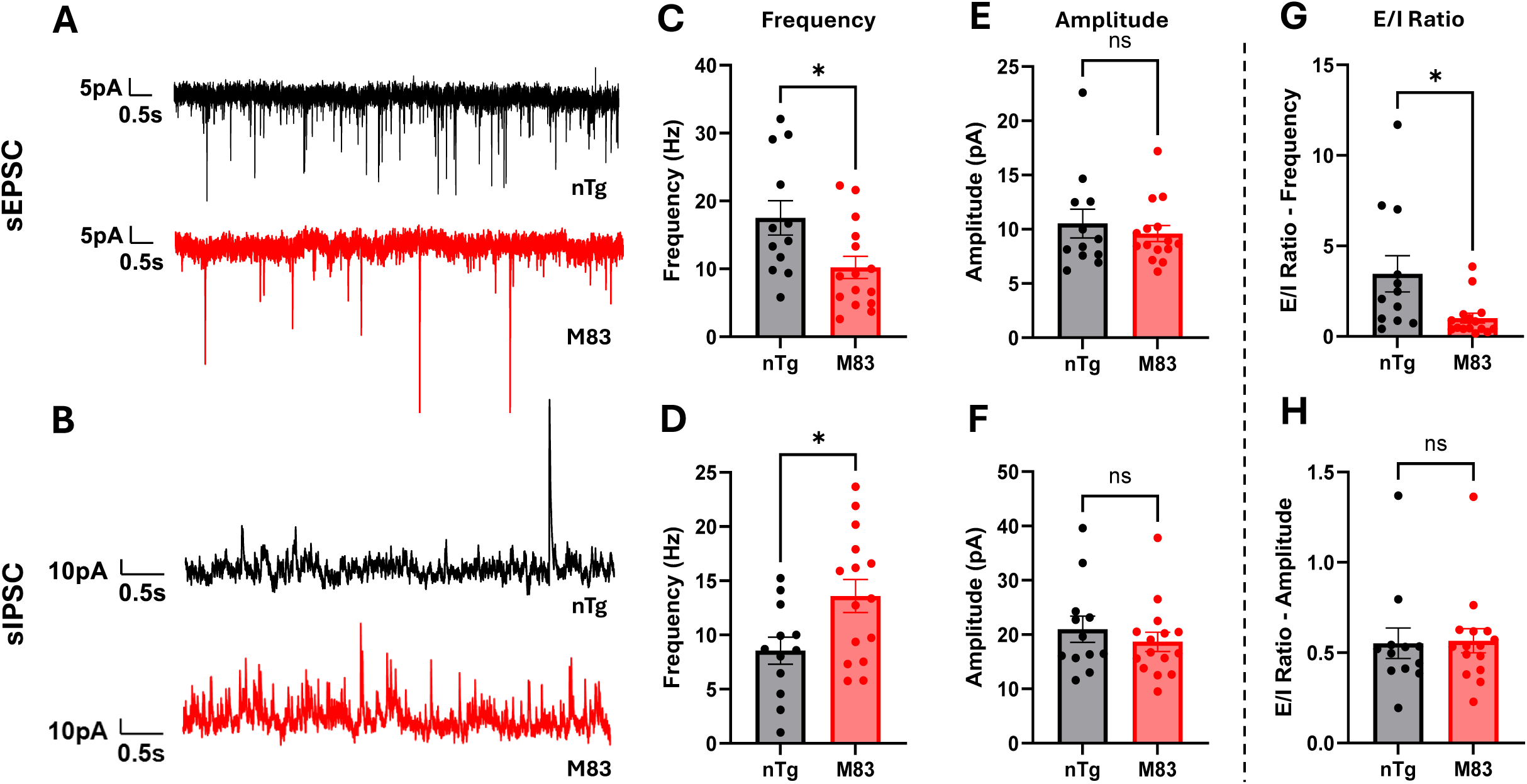
Imbalanced excitatory/inhibitory synaptic transmissions measured in excitatory cortical neurons in M83 mice. **A-B**. Example whole-cell voltage-clamp recordings showing the spontaneous excitatory postsynaptic currents (sEPSCs) and inhibitory postsynaptic currents (sIPSCs) of layer 2/3 motor cortical excitatory neurons from 8–9-month female M83 and age-matched nTg mice. **C-D**. sEPSC frequency of excitatory neurons in M83 mice was significantly decreased compared with nTg mice, while sIPSC frequency of excitatory neurons in M83 mice was significantly increased compared with nTg mice. **E-F**. The mean amplitude of both sEPSC and sIPSC maintained the same between M83 and nTg mice. Results in C - F indicate altered presynaptic excitatory and inhibitory inputs and a preserved postsynaptic response. **G-H**. The sEPSC/sIPSC ratio of the same recorded excitatory neuron was calculated and compared between two genotypes. M83 mice displayed significantly reduced E:I ratio in frequency compared with the nTg group. Suggesting a shift in the E/I balance toward inhibition at the synaptic level. Excitatory neurons: nTg n=12, M83 n=15; Unpaired Mann-Whitney t-test. *: P<0.05.

Immunohistology of M83 mice revealed a preferential colocalization of phosphorylated α-syn aggregates with excitatory neurons rather than with PV interneurons in layer 2/3 of motor cortex (**Fig. 10A-B**). Quantitative analysis confirmed a higher degree of overlap between pathological aggregates and excitatory neuronal populations (**Fig. 10C**). This cell-type-selective distribution indicates that excitatory neurons are more vulnerable to the accumulation of α-syn pathogens.

**Figure 10.**
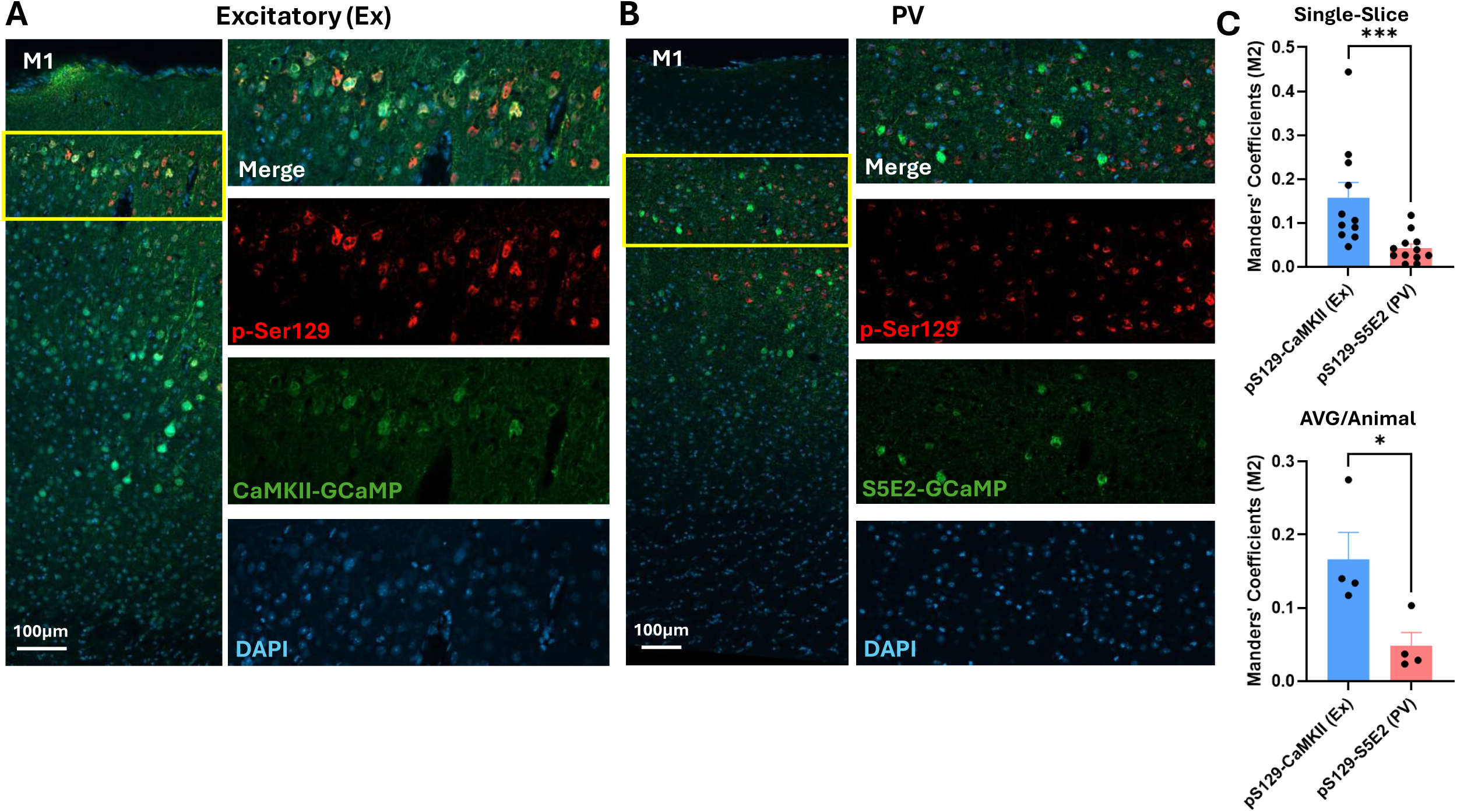
α-syn aggregates preferentially accumulated in excitatory neurons rather than PV neurons in M83 mice. **A-B**. After two-photon imaging, M83 mice brain sections were stained with anti-pSer129 (for phospho-Ser129) and anti-GFP (for GCaMP) antibodies. pSer129 co-localized more prominently with excitatory neurons (CaMKII-GCaMP) than with PV neurons (S5E2-GCaMP). **C**. Colocalization in layer 2/3 of M1 was quantified by Mander’s coefficient. A significantly higher degree of overlap was observed between pSer129 and excitatory neurons (Ex) compared to PV neurons, both in pooled brain slices and averaged value per-animal. Excitatory neurons: n=11 slices of 4 M83 mice; PV neurons: 12 slices of 4 M83 mice. Unpaired Mann-Whitney t-test: *, ***: P<0.05, 0.001.

Together with the electrophysiological findings, these results indicate that altered intrinsic membrane properties of excitatory neurons are induced by selective accumulation of α-syn aggregates, whereas PV neurons as the powerful regulators of excitatory neurons are largely exempted. This selective neuronal vulnerability in motor cortex largely clarifies the mechanisms of the neuronal dysfunction observed in vivo.

## Discussion

This study addresses a critical gap in understanding how α-syn pathology affects cortical function and motor behavior in the early stage of synucleinopathy. This gap is especially important given the growing recognition of cortical pathogenesis in Parkinson’s disease in both clinical and animal studies [13, 39, 40]. Additionally, DLB is defined by cortical synucleinopathy but the mechanism of its neuronal dysfunction and progression are largely unknown. Our findings provide evidence that motor cortical dysfunction is not only present in M83 mice but that it emerges in a cell-type-selective and temporally structured manner during motor learning, suggesting that imbalanced cortical circuits in early stage may initiate and amplify relevant features in disease progression.

One of the key pathological mechanisms underlying synucleinopathy is the toxicity mediated by Ca^2+^ dependent processes. Accumulating evidence from both in vivo and in vitro studies indicates that pathological α-syn interferes with intracellular Ca^2+^ homeostasis and buffering mechanisms, suggesting its early impact on disease progression [17, 41]. Potential mechanisms underlying differential vulnerability have been characterized in the dopaminergic system, where VTA dopaminergic neurons are largely spared from degeneration, in contrast to SNc neurons. Although both SNc and VTA dopaminergic neurons exhibit tonic pacemaker activity associated with substantial metabolic demand, their susceptibility to Ca^2+^-mediated stress differs. SNc neurons use L-type Ca^2+^ channel (LTCC) for pacemaker function but VTA neurons use NaV channels [42, 43]. In addition, VTA neurons possess greater Ca^2+^-buffering capacity than SNc neurons due to their higher expression levels of the Ca^2+^-binding protein calbindin [44].

Similar principles may extend to cortical circuits, where differences in Ca^2+^ buffering ability between excitatory neurons and PV interneurons could contribute to selective vulnerability. In particular, PV acts as a high-affinity, slow Ca^2+^-binding protein that serves as a powerful intracellular Ca^2+^ buffer, limiting excessive Ca^2+^ accumulation and its spatial distribution [45]. This specialized buffering capacity supports the sustained high-frequency tonic firing characteristic of PV interneurons and may protect them from Ca^2+^-mediated stress as observed in this study and other proteinopathies [46]. In contrast, excitatory pyramidal neurons possess comparatively weaker Ca^2+^-buffering capacity, potentially rendering them more susceptible to Ca^2+^ dysregulation. As a result, elevated intracellular Ca^2+^ level could further promote α-syn oligomerization, which can form Ca^2+^-permeable membrane pores [47], as well as regulate abnormalities in α-syn lipid binding and disrupt synaptic vesicle interactions [48]. More comprehensive investigations into related mechanisms are needed at the molecular level and may provide insights into the link between intrinsic neuronal activity, Ca^2+^ homeostasis, and selective vulnerability in the cortex.

Interestingly, motor exercise gradually normalized—at least transiently—the hypoactive state of excitatory neuronal activity, accompanied by a reduction in PV interneuron activity. This may reflect a compensatory mechanism and interactions between excitatory and PV neurons. However, behavioral performance did not improve even at later sections of training, nor was there evidence of population-level plasticity, as reflected by the absence of repeatable sequential activation patterns in excitatory neuronal ensembles. The impaired plasticity of cortical neurons in M83 mice may result from a shift away from the critical network state required for motor learning [49], which enables learning-induced synaptic reorganizations, including synaptic potentiation and depression. Maintenance of this critical state is believed to rely heavily on a precisely balanced E/I network [49, 50]. Notably, this cortical dysfunction was observed at an early stage of disease, preceding overt neurodegeneration and potentially contributing to the emergence of early motor symptoms. These findings may therefore have important clinical implications for early diagnosis and intervention.

It is worth noting that, compared with our finding of reduced excitability in excitatory neurons of M83 mice, previous studies in different synucleinopathy models have reported variable and sometimes controversial alterations in neuronal excitability in vitro. Some studies demonstrated selectively increased excitability, such as in substantia nigra dopaminergic neurons [51], and motor cortical intratelencephalic neurons (ITNs) [19], whereas others reported decreased excitability in substantia nigra pars compacta dopaminergic neurons [52], layer 5 pyramidal cortical neurons [53], and human-induced pluripotent stem cells (hiPSCs) dopaminergic neuron [41]. These discrepancies may arise from differences in disease models and the stages in disease progression. They may also reflect the heterogeneous impact of α-syn pathology across different brain regions and cell types. Therefore, further comparative studies across different models, combined with molecular-level investigations, are needed to better elucidate the mechanisms by which α-syn pathology alters the intrinsic properties of neurons.

The M83 mouse model used in this study recapitulates the chronic progression of synucleinopathy especially in familial cases, through expression of the human A53T mutant form of α-syn, one of the first identified and most common pathogenic mutations associated with familial Parkinson’s disease. Compared with models that overexpress wild-type human α-synuclein, the A53T model helps reduce the ambiguity regarding whether observed phenotypes result merely from elevated levels of physiological membrane-associated α-helical α-synuclein or from increased formation of pathological α-syn species. The homozygous mice used typically develop motor symptoms between 8 and 16 months of age, whereas heterozygous mice develop symptoms after 22 months, providing a long enough time window for investigating early-stage pathophysiological alterations. In addition, due to the characteristics of the prion protein promoter used in the transgene, nigrostriatal dopaminergic neurons are not among the earliest affected neuronal populations, making the M83 line particularly suitable for studying cortical pathology more independently.

The cortical pathophysiological features identified in this study only reflect early-stage dysfunction. As the disease progresses, additional alterations may emerge in cortical PV neurons at later stages, or that subtle changes may already occur during the early stage but remain undetected in the present study. Therefore, it would be worthwhile to further investigate the selective vulnerability and functional properties of PV neurons across different disease stages and synucleinopathy models. Such investigations may provide important insights into pathogenic mechanisms beyond the classical midbrain mechanism and contribute to more comprehensive understanding of motor impairments in synucleinopathy.

## Methods

### Animals

This study used 8-9-month female homozygous transgenic mice (M83, JAX# 004479)) [21], that express mutant human A53T α-synuclein under the direction of the mouse prion protein promoter, and their age matched non-transgenic controls of the same strain background. All mice were housed in animal facilities of New York University Grossman School of Medicine with experimental and housing conditions approved by the Institutional Animal Care and Use Committee (IACUC) and in accordance with National Institutes of Health (NIH) guidelines, which meet or exceed the ARRIVE guidelines (Animal Research: Reporting of In Vivo Experiments).

### Surgical preparation for in vivo imaging

Mice were deeply anesthetized through intraperitoneal injection of ketamine (100mg/kg) - xylazine (10mg/kg xylazine) cocktail. Viral injections targeting layer 2/3 of the caudal forelimb area (CFA) of the mouse primary motor cortex (coordinates: AP 0.3 mm, ML 1.5 mm) were made through a small (∼ 0.5mm in diameter) opening of skull. Each injection lasted 10 minutes through a glass pipette (Sutter Instrument) to deliver 200 nl of the viral solution by a microinjector. Next, a 3 mm diameter craniotomy was carefully made by a dental drill on the skull centered by the viral injection site, followed by the implantation of a glass coverslip (Warner Instruments) onto the exposed dura surface which was glued (Loctite) to the skull. Finally, a custom-designed metal head plate was mounted onto the skull with the dental acrylic cement to enable head fixation during recording. Imaging and behavioral sessions were conducted 2-3 weeks after the surgery.

### Behavioral paradigm and assessment

The running treadmill was custom built on an optical breadboard (Thorlabs) where a polyester belt was driven by a DC motor that was controlled by a single-board microcontroller and a motor shield (Arduino) with custom programs (IDE). Ca^2+^ imaging and behavioral tracking were synchronously recorded by a data acquisition card (National Instruments) with MATLAB programs.

All mice underwent 30 minute of motor training in 6 consecutive sessions, each comprising 5 forced forelimb running trials of 45s in length. This training paradigm was developed based on previous studies [35, 54] and modified to better reflect the gradual progression of learning. Consecutive running trials were programmed to be separated by randomized intertrial intervals of 13–15 s to minimize the effect of time perception and anticipation. Forelimb movements were continuously captured through the infrared camera in the dark over the training process from both front and side views. Continuously captured image series were analyzed through deep learning-based analyzing tool (DeepLabCut) [22], where we manually labeled the paw positions in a small subset of image frames before network training to identify their positions in the remaining images. Then, 3D front paw positions were extracted to calculate gait parameters, including swing time, stride length, step number, and trajectory correlations.

### In vivo two-photon imaging

In vivo Ca^2+^ imaging was conducted on awake mice through commercial two-photon microscopy (Olympus) with a 25x objective (Olympus) at a resolution of 256 x 256 pixels and 2× digital zoom. The mice were head-restrained on the treadmill and remained in a resting state for 10 minutes before training for habituation. Ca^2+^ indicator GCaMP was excited by 920 nm femtosecond laser (Spectra Physics) and emission light was collected through a 525/50nm bandpass filter. Focal plane of each mouse was randomly selected within layer 2/3 of M1. Continuous imaging was performed throughout the entire training process, and mice exhibiting evident focal plane drift were excluded from analysis.

### Two-photon imaging analysis

Two-photon recordings during motor training were preprocessed through Suite2p toolbox [55] to correct evident motions across frames. Neurons with overfilled fluorophore signal or regions of interest (ROIs) that did not match the shape of cell bodies were excluded manually from further analysis, followed by neuron registration, extraction of fluorescent traces and calculation of fluorescent changes relative to baseline (ΔF/F). The frequency, amplitude and area under the curve (AUC) of the Ca^2+^ events were calculated referring to our prior studies [56]. The baseline used in calculating these parameters was determined by the mean activity within a sliding window spanning 10% of recording length with the minimal standard deviation (SD). Excitatory neurons were classified as hypoactive or hyperactive if their activity frequency was lower than 1 or higher than 3 transients/min, respectively. Running activated or suppressed neurons were identified in each training section through paired t-test by comparing averaged neuronal activity during running versus resting periods of all trials within that section, with a significance threshold of p < 0.05. Heatmap was plotted with normalized ΔF/F and neurons were sorted according to their peak time in the first trial of each training stage. To quantify the repeatability of sequential activation, peak-time-sorted activity sequences in each trial were first obtained from the same group of activated neurons, after which pairwise Pearson’s correlation coefficients of the sequences were calculated across trials from the same section.

### Brain slice whole-cell recordings

M83 or nTg female mice at 8–9-months of age were cardiac perfused with oxygenated solution that contained (in mM): 87 NaCl, 2.5 KCl, 1.25 NaH2PO4, 25 NaHCO3, 10 D-glucose, 75 sucrose,1.3 ascorbic acid, 7 MgCl2, and 0.5 CaCl2. Mouse brains were quickly extracted in this ice-cold solution and coronally sectioned into 250 μm slices by a vibratome (Leica) before being transferred to oxygenated (95% O2 / 5% CO2) artificial cerebrospinal fluid (aCSF) (in mM): 124 NaCl, 2.5 KCl, 1.25 NaH2PO4, 26 NaHCO3, 10 glucose, 1.5 MgCl2, and 2.5 CaCl2 at 35°C for 20 min. Then, the brain slices were recovered for 1 h in aCSF at room temperature.

Whole-cell current-clamp recordings were conducted with perfusion of 30°C oxygenated aCSF under 40x objective (Olympus). GCaMP positive cell bodies locate in layer 2/3 of M1 were first identified through FITC channel for specific cell types prior to recording in the differential interference contrast (DIC) channel. Glass pipettes were filled with the current-clamp internal solution (in mM): 127 K-gluconate, 8 KCl, 10 phosphocreatine, 10 HEPES, 4 Mg-ATP, 0.3 Na-GTP or voltage-clamp internal solution (in mM): 130 Cs-methanesulfonate, 4 TEA-Cl, 10 phosphocreatine, 10 HEPES, 1 QX-314, 4 Mg-ATP, 0.3 Na-GTP. For F-I curve and intrinsic firing features, current-clamp recordings were conducted with a series of depolarization current steps injected starting from -80 mV. Data were recorded at a sampling rate of 20 kHz using a MultiClamp 200 amplifier (Molecular Devices). Cells with access resistance exceeding 25 MΩ were excluded from further analyses. For synaptic transmissions including sEPSCs and sIPSCs, voltage-clamp recordings were conducted with membrane potential held at -70 mV or 0 mV respectively. Data was processed by Clampfit (Molecular Devices) followed by analysis through custom MATLAB scripts.

### Western blot

Following in vivo experiments, mouse subjects were cardiac perfused with PBS and one brain hemisphere was homogenized in modified RIPA buffer (50 mM Tris-HCl, 150 mM NaCl, 1 mM EDTA, 1% Nonidet P-40, pH 7.4) with protease and phosphatase inhibitors cocktail (1X protease inhibitor mixture (cOmplete, Roche), 1 mM NaF, 1 mM Na3VO4, 1 nM phenylmethylsulfonyl fluoride (PMSF), and 0.25% sodium deoxycholate). The homogenates were centrifuged at 20,000 x g for 20 min at 4°C. Then the supernatant was collected in pre-chilled tubes as Low-Speed Supernatant (LSS). Protein concentration was measured with the BCA assay and adjusted to the same concentration with the homogenization buffer. To further separate insoluble sarkosyl pellet (SP) and soluble High-Speed Supernatant (HSS), LSS aliquots with equal amount of protein (2 mg) were mixed with 10% sarkosyl in RIPA buffer to achieve 1% sarkosyl final concentration, before ultra centrifugation (Optima MAX-XP, Beckman) at 100,000 x g at 20°C for 1 h. The pellets were washed with 1% sarkosyl solution and centrifuged again at the same speed before air dried as SP. All samples were diluted by modified O+ buffer (62.5 mM Tris-HCl, 10% glycerol, 5% β-mercaptoethanol, 2.3% SDS, 1 mM EDTA, 1 mM EGTA, 1 mM NaF, 1 mM Na3VO4, 1 nM PMSF, and 1X protease inhibitor mixture) and electrophoresed on 12% SDS-PAGE gels. After protein transfer, membranes were fixed with 4% paraformaldehyde (PFA) for 30 min at room temperature. The membranes were blocked with 5% milk in Tris buffered saline with 1% TritonX-100 (TBST) for 1 h before incubation with combinations of following primary antibodies: Phospho-α-syn (pSer129) rabbit antibody (1:2000, D1R1R clone, Cell Signaling Technology), αSyn 211 mouse antibody (1:2000, Thermo Fisher), and GAPDH rabbit antibody (1:4000, Cell Signaling Technology) overnight at 4°C. Secondary antibodies (goat anti-rabbit IRDye® 800CW and goat anti-mouse IRDye® 680RD 1:10,000, LI-COR Biosciences) were applied consequently followed by image scanning by Odyssey CLx scanner (LICOR Biosciences) and quantified by LI-COR Image Studio Lite 5.2.

### Immunohistochemistry

After in vivo tests, a mouse was transcardiac perfused with PBS and one hemisphere of the subject mice was fixed in 4% paraformaldehyde at 4°C overnight and then immersed in 30% sucrose in PBS at 4°C for cryoprotection before freezing it in OCT. Then, the frozen brain was coronally sectioned into 30 - 40 μm sections using cryostat and stored in cryoprotectant (30% ethylene glycol, 30% sucrose in 0.1 M phosphate buffer, pH 7.4) at -20°C before use. Free floating brain slices were washed in PBS before 2 h room temperature blocking in PBS solution containing 3% normal goat serum, 2% bovine serum albumin and 0.5% Triton X-100. Slices were then incubated with combinations of following primary antibodies overnight at 4°C: Phospho-α-syn (pSer129) rabbit antibody (1:1000, D1R1R clone, Cell Signaling Technology), GFP chicken antibody (1:1000 Abcam), Tyrosine hydroxylase rabbit antibody (1:1000, Millipore), NeuN mouse antibody (1:1000, Millipore). The brain sections were then washed with PBS and incubated with combinations of following secondary antibodies (1:1000) at room temperature for 2 h: goat anti-rabbit Alexa Fluor 568/488, goat anti-mouse Alexa Fluor 488, and goat anti-chicken Alexa Fluor 488 (ThermoFisher). To stain the nuclei, DAPI (1:2000) was applied for 20 min at room temperature after washing of the secondary antibody. After extra washing steps with PBS, the sections were mounted in ProLong Gold antifade reagent (ThermoFisher) before imaging through an LSM 800 Zeiss confocal microscope with 10× or 20x objectives. Quantification of confocal images were performed by Fiji software (ImageJ).

### Statistics

Statistical analyzing of both behavioral performance and neuron activity were accomplished by GraphPad Prism. Paired or unpaired Mann-Whitney or parametric two-tailed t-test were used following Shapiro–Wilk normality test. Two-way ANOVA were used followed by Tukey’s multiple comparisons test or Fisher’s LSD multiple comparison test, with the specific test for each figure are listed in the figure legends.

## Author Contributions

X.Y. and E.M.S. conceived the project and wrote the article. X.Y. performed most of the experiments and data analyses. C.J. and S.S. provided guidance on electrophysiological recording and data analyses. N.H. contributed to the testing and adaptation of the DeepLabCut toolbox. Y.L. maintained the animal colonies. E.M.S. supervised the project.

## Acknowledgments

This work was supported in part by R56 AG083436, R01 AG032611, R01 NS077239, R01 NS120488 (E.M.S.), and NIA K01 AG092910 (C.J.). We also thank New York University (NYU) Langone Health’s Ion Laboratory (RRID: SCR_021754) and Microscopy Laboratory (RRID: SCR_017934) for experimental and technical support.

## Availability of Supporting Data

The experimental dataset presented and analyzed in this study are available from the corresponding author upon a reasonable request.

